# A method for estimating dynamic functional network connectivity gradients (dFNG) from ICA captures smooth inter-network modulation

**DOI:** 10.1101/2024.03.06.583731

**Authors:** Najme Soleimani, Armin Iraji, Theo G.M. van Erp, Aysenil Belger, Vince D. Calhoun

**Affiliations:** Tri-Institutional Center for Translational Research in Neuroimaging and Data Science (TReNDS), Georgia State University, Georgia Institute of Technology, and Emory University, Atlanta, Georgia, USA; Clinical Translational Neuroscience Laboratory, Department of Psychiatry and Human Behavior, UC Irvine, Irvine, California, USA; Department of Psychiatry, University of North Carolina, Chapel Hill, North Carolina, USA

**Keywords:** Dynamic functional network connectivity (dFNC), Gradient, Dynamic functional network connectivity gradient (dFNG), Independent Component Analysis (ICA), Schizophrenia

## Abstract

Dynamic functional network connectivity (dFNC) analysis is a widely used approach for studying brain function and offering insight into how brain networks evolve over time. Typically, dFNC studies utilized fixed spatial maps and evaluate transient changes in coupling among time courses estimated from independent component analysis (ICA). This manuscript presents a complementary approach that relaxes this assumption by spatially reordering the components dynamically at each timepoint to optimize for a smooth gradient in the FNC (i.e., a smooth gradient among ICA connectivity values). Several methods are presented to summarize dynamic FNC gradients (dFNGs) over time, starting with static FNC gradients (sFNGs), then exploring the reordering properties as well as the dynamics of the gradients themselves. We then apply this approach to a dataset of schizophrenia (SZ) patients and healthy controls (HC). Functional dysconnectivity between different brain regions has been reported in schizophrenia, yet the neural mechanisms behind it remain elusive. Using resting state fMRI and ICA on a dataset consisting of 151 schizophrenia patients and 160 age and gender-matched healthy controls, we extracted 53 intrinsic connectivity networks (ICNs) for each subject using a fully automated spatially constrained ICA approach. We develop several summaries of our functional network connectivity gradient analysis, both in a static sense, computed as the Pearson correlation coefficient between full time series, and a dynamic sense, computed using a sliding window approach followed by reordering based on the computed gradient, and evaluate group differences. Static connectivity analysis revealed significantly stronger connectivity between subcortical (SC), auditory (AUD) and visual (VIS) networks in patients, as well as hypoconnectivity in sensorimotor (SM) network relative to controls. sFNG analysis highlighted distinctive clustering patterns in patients and HCs along cognitive control (CC)/ default mode network (DMN), as well as SC/ AUD/ SM/ cerebellar (CB), and VIS gradients. Furthermore, we observed significant differences in the sFNGs between groups in SC and CB domains. dFNG analysis suggested that SZ patients spend significantly more time in a SC/ CB state based on the first gradient, while HCs favor the SM/DMN state. For the second gradient, however, patients exhibited significantly higher activity in CB domains, contrasting with HCs’ DMN engagement. The gradient synchrony analysis conveyed more shifts between SM/ SC networks and transmodal CC/ DMN networks in patients. In addition, the dFNG coupling revealed distinct connectivity patterns between SC, SM and CB domains in SZ patients compared to HCs. To recap, our results advance our understanding of brain network modulation by examining smooth connectivity trajectories. This provides a more complete spatiotemporal summary of the data, contributing to the growing body of current literature regarding the functional dysconnectivity in schizophrenia patients. By employing dFNG, we highlight a new perspective to capture large scale fluctuations across the brain while maintaining the convenience of brain networks and low dimensional summary measures.

## 1. Introduction

Functional connectivity (FC) refers to the functional coactivation of brain activity between spatially segregated brain regions regardless of their apparent physical connectedness [1]. FC is most often measured during resting state fMRI as a statistical relationship (e.g., correlation) based on temporal similarities to study functional brain networks [1],[2],[3]. Building on the same concept, functional network connectivity (FNC), refers to the interaction between spatially separable, overlapping, temporally coherent brain networks (also known as intrinsic connectivity networks or ICNs) [4]. Traditionally, functional connectivity assumes a constant connectivity pattern over the data acquisition time period [5]. However, dynamic functional connectivity analysis has shown that far from being static, the functional networks captured with fMRI reveal brain fluctuations on the scale of seconds to minutes. These changes are often summarized as movements from one short term state to another, rather than continuous shifts [5], though such measures can also be easily represented via smoothly varying transitions [6], or as overlapping dynamic movies [7]. Dynamic functional connectivity has also demonstrated that the blood oxygenation level dependent (BOLD) signals measured during resting state include important spatio-temporal dynamic properties [8],[9]. Many studies have replicated such reproducible patterns of network activity that move throughout the brain [5], [9],[10].

The emergence of dynamic functional connectivity has revolutionized our ability to study underlying brain systems by providing information about the temporal changes in brain connectivity and various types of brain dynamic properties [9]. There has been a growing interest in studying the temporal reconfiguration of brain functional connectivity suggesting that the spatial and temporal properties of neural activity interact through several spatiotemporal scales [11],[12]. The spatial dynamics of the brain constitute a multifaceted domain of inquiry within neuroscience. These dynamics pertain to the intricate patterns of functional connectivity and organization that underlie cognitive processes and behaviors [9],[13]. Functional networks, composed of spatially distributed brain regions, form the basis of these dynamics, and their organization reflects the underlying neural architecture [13]. Understanding the spatial dynamics of the brain is pivotal not only for advancing our fundamental knowledge of brain function but also for elucidating the pathophysiology of neurological and psychiatric conditions.

Alongside these endeavors, recent years have witnessed empirical studies focused on a novel approach investigating the temporally static spatial topography of brain connectivity known as spatial gradients [14]. Recent research has also underscored the importance of cortical gradients [14], which reveal smooth transitions in connectivity patterns across the cortex. These gradients provide valuable insights into the spatial organization of the brain’s functional networks, shedding light on their interplay and facilitating the identification of individual differences and alterations associated with neurological disorders [15],[16]. Adopting a macroscale perspective on cortical organization has already provided insights into how cortex-wide patterns relate to cortical dynamics [17].

Building upon this understanding, we propose two innovative approaches in our study. Firstly, we introduce subject-specific reordering of independent component analysis (ICA) networks (i.e., ICNs) based on the inter-component functional connectivity gradient (i.e., FNG). Cortical gradients help us understand the spatial organization of functional connectivity patterns across the brain. The use of low-dimensional representations of functional connectivity provides a unified perspective to efficiently explain core organizing properties of the human cerebral cortex, linking specific regions, networks, and functions. Cortical gradients establish a framework to study brain organization and the covariation between spatial and temporal factors [15] through quantifying topographic principles of macroscale organization [18], thus providing insight into brain organization in healthy individuals as well as in individuals with mental disorders [19]. By leveraging higher order statistics and spatial constraints, we automatically separate canonical networks and subsequently re-order them based on smooth gradients at the individual subject level. To put it simply, we initially identify the well-known brain networks, and then, we organize them in a way that ensures a seamless and gradual transition between these networks using gradients, all while considering the unique connectivity patterns of each individual subject’s brain. This process allows us to create a more personalized and precise understanding of how these brain networks function in each person, considering individual variability. These two strategies are complementary, as ICA naturally identifies and separates reliable and replicable overlapping spatial networks (regardless of their topological smoothness) using higher order statistics, whereas gradient approaches focus on smoothly varying patterns which are typically orthogonal and ordered by variance. This joint approach enables a more spatially precise and personalized characterization of brain connectivity patterns while also leveraging higher order statistics in the original network determination.

Secondly, we present the novel concept of dynamic gradient reordering, recognizing the need to study how the brain’s functional organization changes over time. Most dynamic functional network studies assume fixed spatial maps and evaluate transient changes in coupling among independent component time courses [4],[20]. In contrast, cortical gradients offer an insight into the smooth and continuous transition of states across the brain by representing brain connectivity in a continuous, low-dimensional space to identify the brain functional hierarchies. Furthermore, they identify spatially distributed patterns of connectivity which reflect the underlying architecture of the brain [14], and how it dynamically reconfigures in response to different cognitive process, suggesting that the temporal dynamics tend to be shaped by the functional geometry [21]. By examining the dynamic nature of cortical gradients, we aim to open a window into the temporal dynamics of atypical macroscale organization across clinical conditions and provide insights into the flexibility and adaptability of brain networks. These innovations can potentially pave the way for a comprehensive investigation of the spatiotemporal organization of the human brain, offering a deeper understanding of its functional dynamics and potential implications for various neurological and psychiatric conditions.

In addition, incorporating both spatial and temporal properties into the summarization step of functional connectivity analysis can be especially important in the context of complex mental illnesses such as schizophrenia since the dynamic nature of brain disruptions can be captured, accounting for inter-individual variability, and monitoring of treatment responses, thus offering comprehensive insights into the disorder’s pathophysiology and potential biomarkers. Schizophrenia is one of the most debilitating psychiatric disorders characterized by hallucinations, delusions, and disordered thinking [22],[23],[24]. A growing body of evidence supports alterations in functional connectivity within and between brain networks associated with the illness [20],[23],[25].

Traditional static functional connectivity analyses using task-based and resting-state functional magnetic resonance imaging have provided valuable insights into the aberrant connectivity associated with schizophrenia [26],[27]. These studies have identified disrupted connectivity within and between different functional networks, including the default mode network, salience network and executive control network [28]. However, these approaches have used static functional connectivity, ignoring different states of brain dynamics. Furthermore, the alterations in task-related connectivity are often related to impaired task performance since it presents a challenge in interpreting the nature of the relationship between altered brain connectivity and impaired task performance. For instance, it may be unclear whether the observed alterations in connectivity are specific to the task being performed or are indicative of a more generalized disruption in brain function.

Previous approaches to resting state dynamic functional connectivity in schizophrenia have shown significantly stronger connectivity between the thalamus and sensory network, and reduced connectivity between putamen and sensory network [20],[24],[4]. However, most existing studies regarding schizophrenia focus on specific brain regions or networks rather than the whole brain or disregard the dynamic properties of the brain.

In the context of schizophrenia, applying gradient-based approaches to dynamic functional network connectivity analysis holds great potential for expanding our understanding of the disorder [19]. By exploring the dynamic functional network connectivity gradients (dFNGs), we aim to uncover links between schizophrenia and the hierarchical organization and transition of functional brain networks.

In this study we propose a novel approach, which aims to investigate the temporal dynamics of functional network connectivity gradients, and we explore the alterations in connectivity gradients in a group of individuals with schizophrenia (SZ) in comparison with healthy controls (HC) matched with the patients in terms of mean age and gender distribution. To our knowledge, no fMRI studies have focused on the dynamics of the whole brain organization using a gradient-based approach, nor have they combined smoothly varying gradients with whole brain networks defined via ICA. To this end, we first compute the reordered FNC in static and dynamic sense, ordered by the variance explained in the initial functional connectivity matrix. The proposed method then uses *k*-means clustering to cluster the reordered dFNG into a set of distinct states, and computes several global dynamic metrics that are compared between groups.

## 2. Materials and Methods

### 2.1. Participants

We evaluate our framework on resting-state functional magnetic resonance imaging (rs-fMRI) data from a cohort consisting of 160 healthy controls (115 males, 45 females, mean age 37.03) and 151 individuals with schizophrenia (115 males, 36 females, mean age 38.76) with similar mean age and gender distribution. The subjects were recruited across seven different sites in the United States as a part of the Functional Imaging Biomedical Informatics Research Network. All patients included in the study had been diagnosed with schizophrenia based on the Structured Clinical Interview for DSM-IV-TR Axis I Disorders (SCID-I/P). Exclusion criteria for both schizophrenia patients and healthy volunteers included a history of major medical illness, contraindications for MRI, poor vision even with MRI compatible corrective lenses, an IQ less than 75, a history of drug dependence in the last five years, or a current substance abuse disorder. Patients with extrapyramidal symptoms and healthy volunteers with a current or past history of major neurological or psychiatric illness (SCID-I/NP) or with a first-degree relative with Axis-I psychotic disorder diagnosis were also excluded. All the participants provided written informed consent prior to scanning in accordance with the Internal Review Boards of corresponding institutions (University of California Irvine, University of California Los Angeles, University of California San Francisco, Duke University/University of North Carolina, University of New Mexico, University of Iowa, and University of Minnesota).

### 2.2. Neuroimaging Data and Preprocessing

Imaging data were acquired on a Siemens Tim Trio 3T scanner at six, and on a 3T General Electric Discovery MR750 scanner at one of the seven sites. A total of 162 volumes of BOLD rs-fMRI were collected using echo planner imaging sequences (TR/TE = 2s/ 30ms, FOV = 220 mm, FA = 77°, 32 sequential ascending axial slices of 4 mm thickness with 1 mm gap). All participants were instructed to keep their eyes closed during the scanning.

rs-fMRI data were preprocessed using the statistical parametric mapping (SPM12, http://www.fil.ion.ucl.ac.uk/spm/) toolbox within Matlab 2020b. The first five scans were removed for the signal equilibrium and participants’ adaptation to the scanner’s noise. We performed rigid body motion correction using the toolbox in SPM to correct subject head motion, followed by the slice-timing correction to account for timing differences in slice acquisition. The rs-fMRI data were subsequently warped into the standard Montreal Neurological Institute (MNI) space using an echo-planar imaging (EPI) template and were slightly resampled to 3 × 3 × 3 mm isotropic voxels. The resampled fMRI images were further smoothed using a Gaussian kernel with a full width at half maximum (FWHM = 6 mm).

### 2.3. Spatially Constrained Independent Component Analysis

Independent component analysis (ICA) is a data-driven method capable of recovering a set of maximally independent sources from multivariate data [29]. ICA is a widely-used exploratory tool to study functional brain networks [29]. However, one of the challenges of the standard ICA approach is “order ambiguity”, which indicates that the order of the independent components (ICs) estimated by the standard ICA is arbitrary [30]. Additional prior information can contribute to the solution to this problem. Spatially constrained independent component analysis uses anatomical priors or templates to extract functional brain networks that are similar predetermined templates and maximally independent [31]. Spatially constrained ICA is thus a hybrid approach which allows individual subject ICA analysis while also providing component ordering and correspondence among subjects. This approach leverages the inherent spatial information to guide the decomposition of functional data into meaningful spatially coherent components [31], [32].

After data preprocessing, the functional data for both control and patient groups were analyzed using spatially constrained ICA employing the Neuromark fMRI 1.0 template as implemented in the GIFT toolbox (http://trendscenter.org/software/gift) [8], resulting in 53 intrinsic connectivity networks (ICNs). The Neuromark fMRI 1.0 is an automatic ICA-based template (downloadable from http://trendscenter.org/data) that enables estimation of brain functional networks from functional magnetic resonance imaging to identify reproducible fMRI markers of brain disorders [33]. The ICNs are partitioned into seven subcategories: subcortical (SC), auditory (AUD), visual (VIS), sensorimotor (SM), cognitive control (CC), default mode network (DMN) and cerebellar (CB) components (see Figure 1).

**Figure 1.**
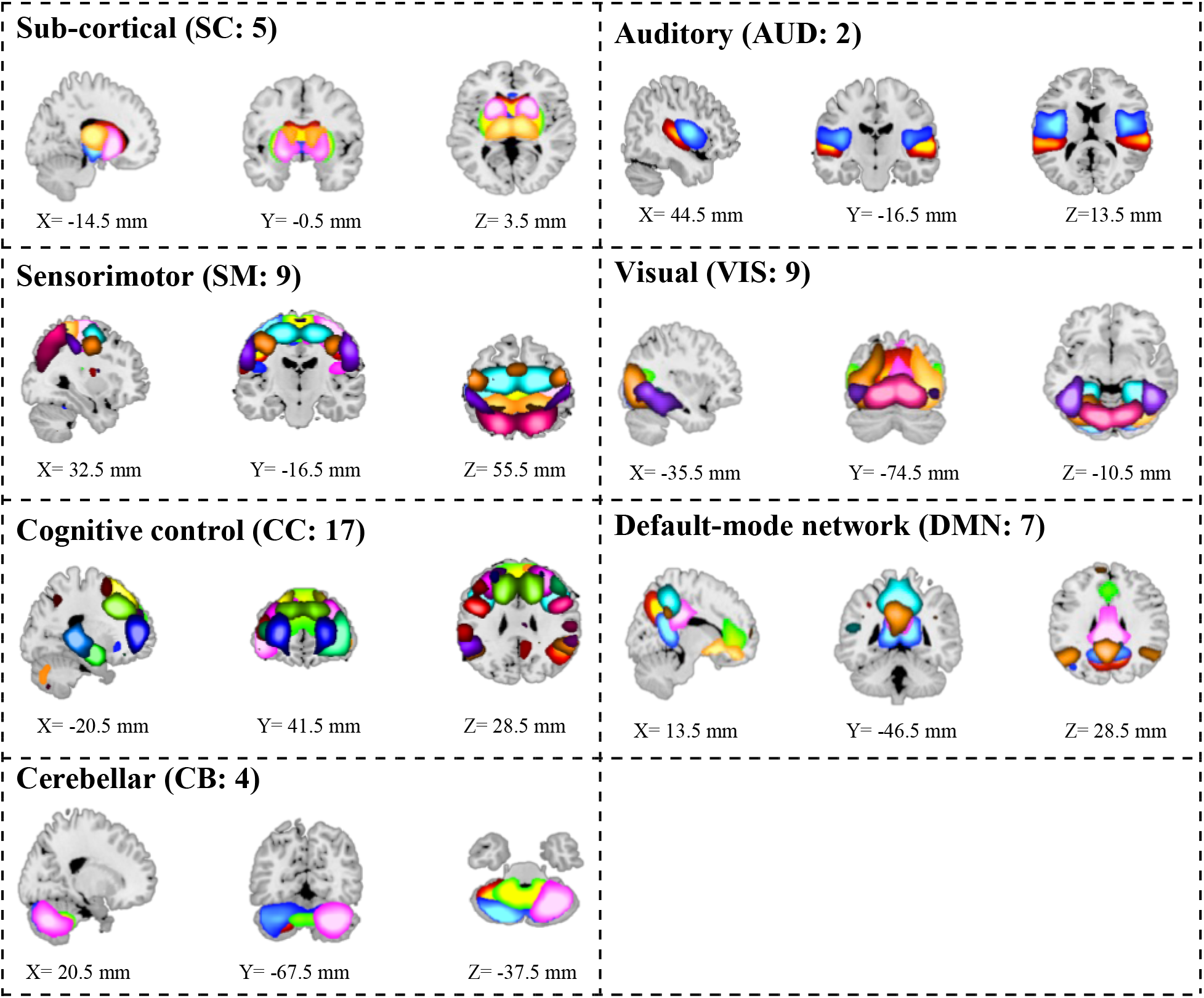
Composite maps of the 53 identified intrinsic connectivity networks (ICNs), divided into seven functional domains. subcortical (SC), auditory (AUD), sensorimotor (SM), visual (VIS), cognitive control (CC), default mode network (DMN) and cerebellar (CB) network.

### 2.4. Functional Network Connectivity Gradients

We computed the static functional network connectivity (sFNC), described as the covariation between ICN full timeseries for each subject, resulting in a 53×53 matrix. Gradients along the sFNC space were computed using the diffusion map approach [18] implemented within the BrainSpace toolbox (https://brainspace.readthedocs.io/en/latest/), which generates efficient representation of complex geometric structures [34], followed by resorting the matrix based on its gradient value. A gradient is an axis of variance along which areas fall in a spatially continuous order [18]. Areas that resemble each other with respect to the feature of interest occupy similar positions along the gradient [35]. Using a diffusion map embedding algorithm that reduces data dimensionality through the nonlinear projection of the vertices into an embedding space, we identified gradient components, estimating the low-dimensional embedding from the high-dimensional connectivity matrix.

Recent empirical studies propose a non-static nature of functional connectivity among different brain regions [9]. To date, the most widely used strategy for examining dynamics in resting state functional network connectivity has been a sliding window approach [8],[5],[10]. This approach involves dividing a continuous timeseries of brain activity into overlapping or non-overlapping windows of fixed duration. By sliding the window along the timeseries, functional connectivity can be computed within each window, capturing the temporal evolution of brain dynamics [8],[10],[36]. Windowed functional network connectivity (windowed-FNC) is computed for each subject using a sliding window approach with a window size of 44 seconds (22 TRs) and strides of 2 seconds (1 TR) [10]. Similar to the static analysis, cortical gradients were computed for each windowed-FNC using the BrainSpace toolbox and reordered subsequently using the diffusion map associated with each time window.

Furthermore, we also developed an approach to track the reordering trajectory, allowing us to create an inter-component ordering synchrony associated with each component for each subject. In our analytical procedure, we begin by generating a sort order matrix for each subject, providing information on the ordering of independent component networks (ICNs), capturing the temporal dynamics of ICN reordering. To assess the level of synchronization across these dynamic changes, we compute cross-correlations across all time lags, followed by extracting the maximum of all lags. This extracted value is subjected to a comparative analysis, allowing us to discern potential differences in the temporal reconfiguration of ICNs between patients and control subjects.

### 2.5. Clustering and Dynamic Functional Network Connectivity Gradient Measures

We used *k*-means algorithm to cluster the dFNG timepoints, partitioning the data into five distinct clusters. The optimal number of clusters was estimated using the elbow criterion, consisting of computing the explained variance as a function of the number of clusters and picking the elbow of the curve [20]. The whole procedure is depicted in Figure 2. *k*-means clustering is a widely used unsupervised algorithm, aimed to partition a given dataset into *k* distinct clusters based on the similarity of data points. The core concept of *k*-means clustering involves finding the centroids of *k* clusters and assigning the data points to the nearest centroid.

**Figure 2.**
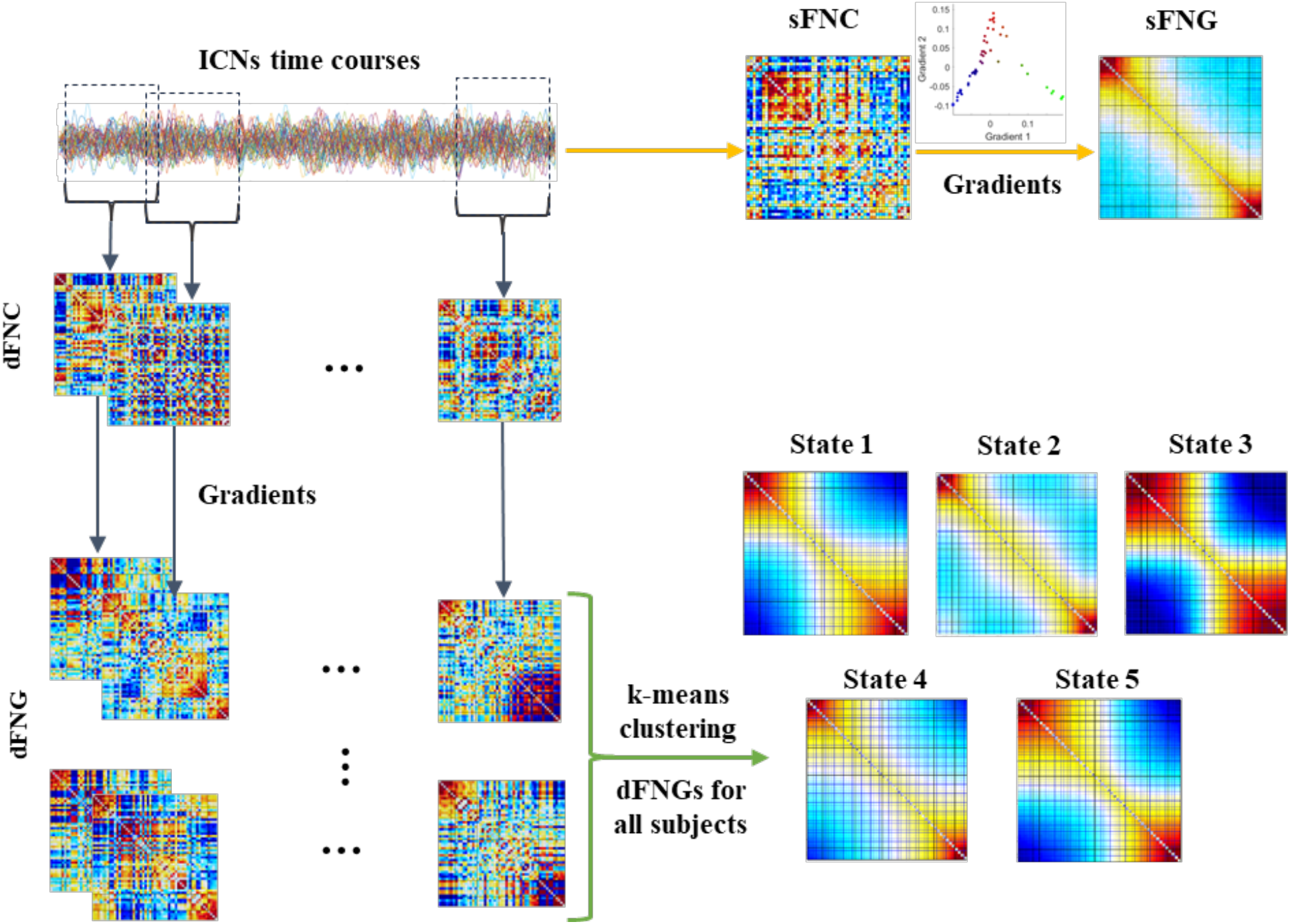
Schematic depicting the proposed method. The fMRI data was preprocessed using standard procedures, and then spatially constrained ICA was run on the data using the Neuromark fMRI 1.0 template, resulting in 53 ICNs. Next, FNC was calculated using sliding window approach. A diffusion map (gradients) was computed for each windowed-dFNC. Each dFNC matrix was reordered based on its gradient, followed by k-means clustering of the reordered dFNC. This resulted in 5 dynamic FNC gradients (dFNGs).

Each FNC gradient represents a weighted combination of the component maps; however, the computed gradients should be corrected for sign ambiguity, since the gradients computed separately from different individuals may not be directly comparable due to sign ambiguity of the eigenvectors. To this end, we utilize group-average gradient matrix as a reference and reverse the sign of each gradient to induce positive correlation before applying clustering analysis. To visualize the weighted combination of the component maps, a spatial map for static functional connectivity gradient (sFNG) was created by thresholding and normalizing each component map (i.e., the largest voxel value equal to one), multiplied by its sign-corrected gradient value and summing them together. We then repeat for all windows to create the dFNG spatial maps. Using the gradient vectors associated with each time point and each subject as the input to k-means clustering, we identified clusters with similar sorting profiles and used the normalized cluster centroids as the weight for the component maps to create spatial maps.

Complementary to examining dynamic changes in connectivity patterns, typical dynamic summary measures [8] such as occupancy, dwell time, and periodicity were calculated to capture the key aspects of dFNG. *Occupancy* refers to the amount of time a brain network spends in a particular state or configuration over the course of a given period, quantifying the number of timepoints each subject spends in each state, providing insight into the stability of a functional state. *Dwell time* refers to the duration or amount of time that a brain network remains in a specific state or configuration before transitioning to another state, reflecting the temporal persistence of a particular functional state. *Periodicity*, however, allows us to assess the oscillatory behavior across brain states.

## 3. Results

We propose a novel approach to leverage the use of higher-order statistics to capture brain networks, coupled with the calculation of gradients to identify a network ordering which maximizes the smoothness in the connectivity. This is then extended to a dynamic connectivity approach, capturing the changes in connectivity over time. We also propose several summary measures and compare these between schizophrenia patients and healthy controls.

### 3.1. Group Differences in static Functional Network Connectivity Gradient (sFNG)

After computing the sFNC for each subject, defined as the temporal correlation between ICNs full time courses, as well as the sFNG, the average sFNC and sFNG for 151 schizophrenia patients (SZ) and that of 160 healthy controls (HC) are computed. Differences in sFNC and sFNG between schizophrenia patients (SZ) and healthy controls (HC) were assessed via two-sample t-test. Figure 3 is illustrative of the average of original sFNC for all subjects (a) and the average of reordered FNC based on gradient 1 (b) and gradient 2 (c).

**Figure 3.**
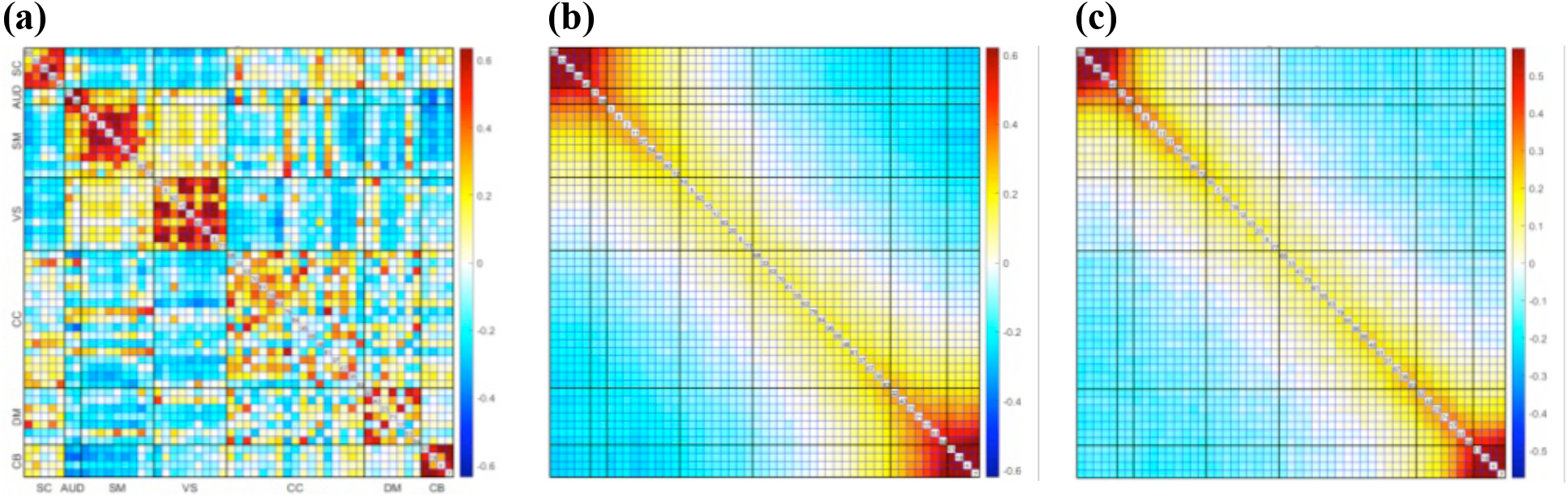
The average of a) original static FNC, b) reordered FNC based on gradient 1, and c) reordered FNC based on gradient 2.

The average of the sign-corrected cortical gradients was computed for patient (SZ) and control (HC) groups and plotted in 2D space and assigning them color. These colors can be informative about the multidimensional interaction between gradients. For HC the three lines correspond to VIS (green), SC/AUD/SM/CC/DMN/CB (blue) and CC/DMN (red) networks. Regarding the SZ, the three lines correspond to VIS (green), SC/AUD/SM/CB (blue) and CC/DMN (red) networks. Figure 4 provides information about the first two cortical gradient interactions.

**Figure 4.**
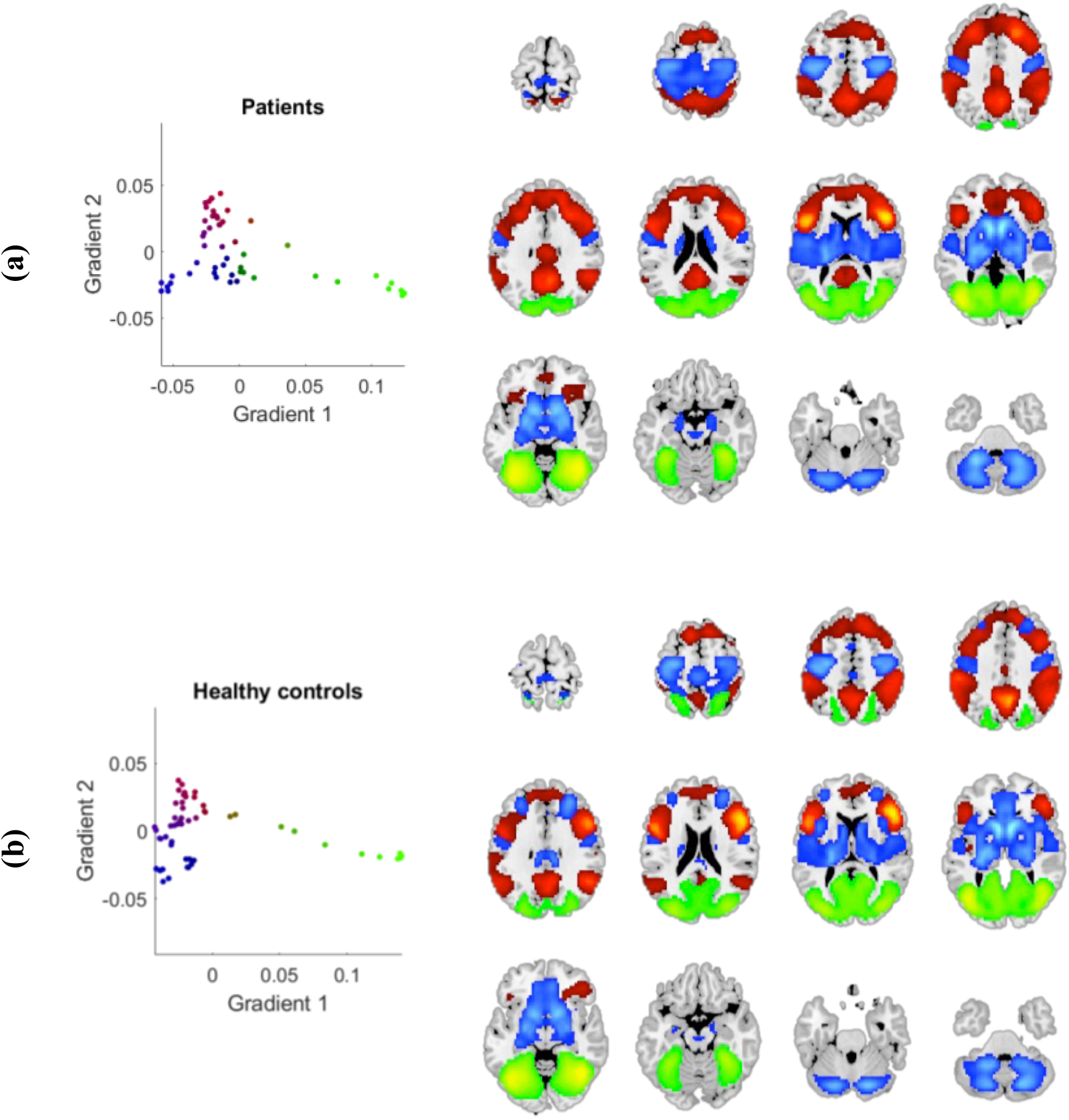
Visualization of the interaction between the average of the first two cortical gradients for a) patients (SZ) and b) control (HC) groups in 2D view. The three dotted patterns correspond to VIS (green), SC/AUD/SM/CC/DMN/CB (blue) and CC/DMN (red) networks for HC and VIS (green), SC/AUD/SM/CB (blue) and CC/DMN (red) networks for SZ.

Regarding the sFNC analysis, compared to the HC, the SZ group showed significantly stronger connectivity between SC, VIS, and SM networks, and significantly weaker connectivity between AUD, SM and VIS networks. As depicted in Figure 5, the schizophrenia patients (SZ) showed significantly weaker connectivity in the subcortical (SC) and cerebellar (CB) domains when compared to controls.

**Figure 5.**
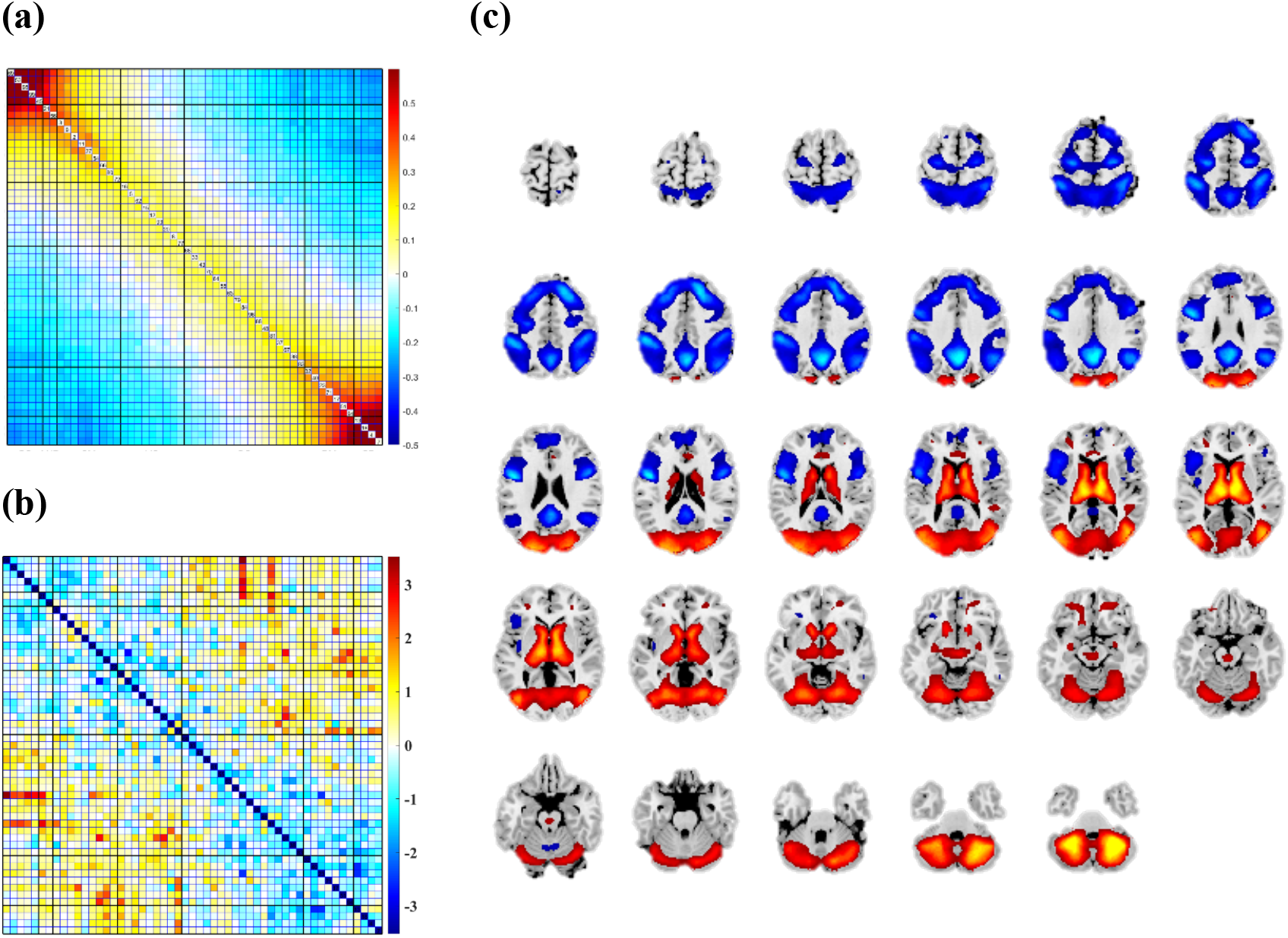
a) The average reordered FNC based on the first gradient associated with schizophrenia patients (SZ), b) the group differences between schizophrenia patients (SZ) and healthy controls (HC) defined as −10 log!“(*pvalue*) × *sign*(*tvalue*), and c) the spatial map associated with the difference between HC and SZ. Regarding the sFNG analysis, the SZ group showed hypoconnectivity in subcortical (SC) and cerebellar (CB) domains.

### 3.2. Group Differences in dynamic Functional Network Gradients (dFNG)

Figure 6 represents the *k*-means cluster centroids associated with dFNC, and dFNGs based on the first and second gradient. A two-sample t-test was applied to investigate the difference in occupancy and dwell time of each state.

**Figure 6.**
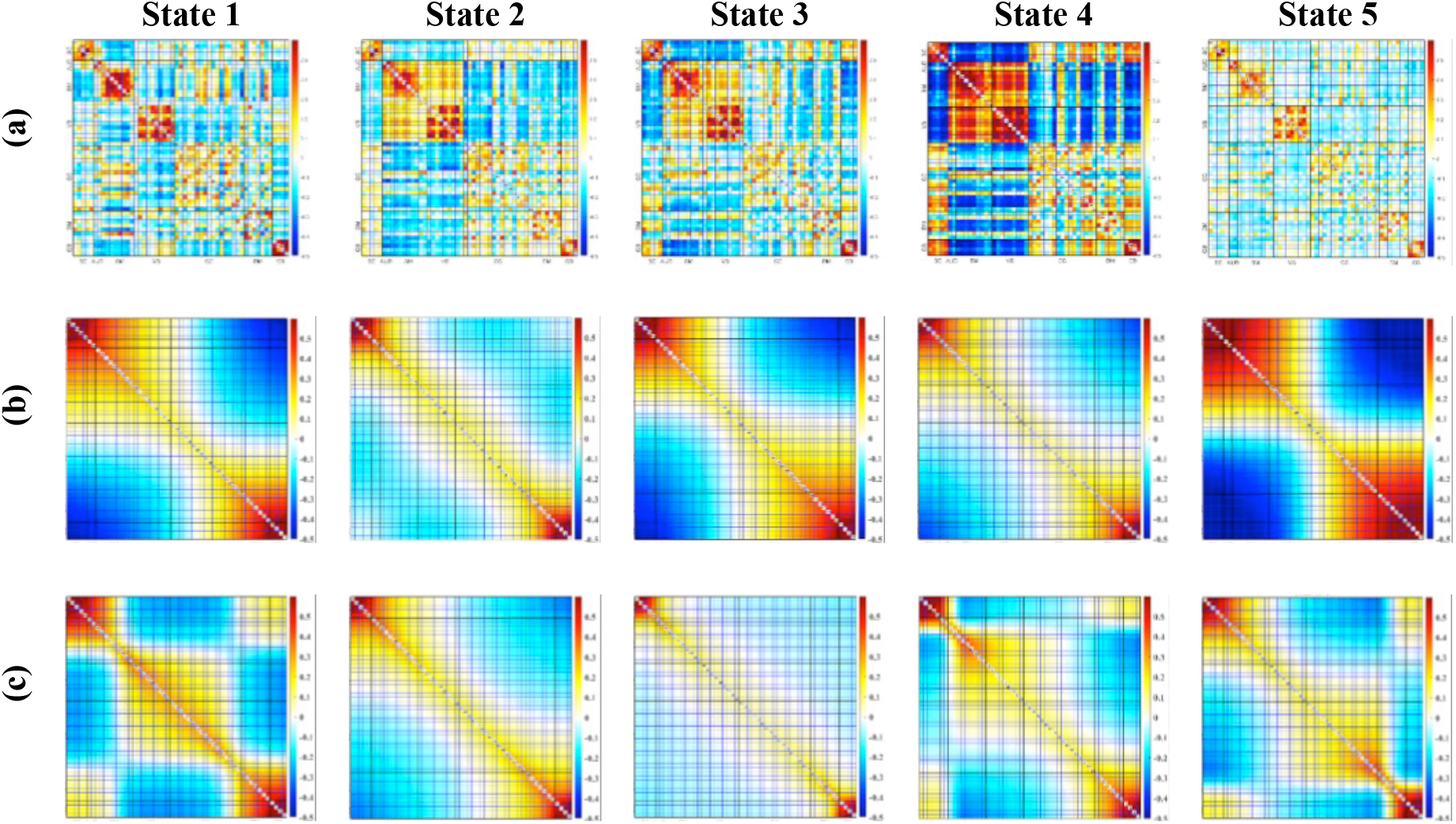
Schematic depicting the state transition (cluster centroids) for a) original dFNC, b) dFNG based on gradient #1, and c) dFNG based on gradient #2.

As is evident in Tables 1 and 2, regarding the dFNG based on the first gradient, the patients (SZ) tend to spend significantly higher duration in state 4 (CB), yet the HCs show a significantly higher occupancy and dwell time in state 3 (SM). However, the second gradient results showed a significantly higher occupancy of the HC group in state 5 (DMN), whereas the SZs spent significantly longer duration in state 1 (CB). All significant results are shown in bold, with those survived after FDR correction are identified with an asterisk.

**Table 1.**
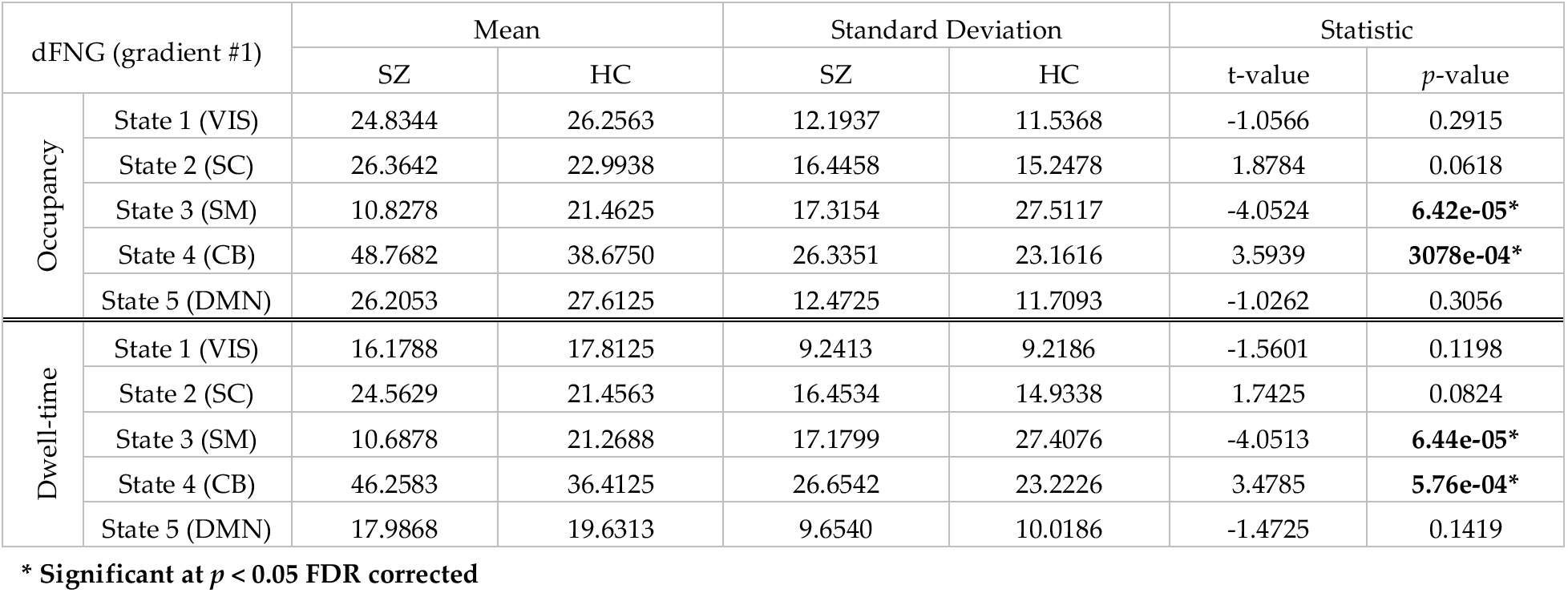
Statistical Results associated with dFNG based on gradient #1.

| dFNG (gradient #1) |  | Mean |  | Standard Deviation |  | Statistic |  |
| --- | --- | --- | --- | --- | --- | --- | --- |
|  |  | SZ | HC | SZ | HC | t-value | p-value |
| Occupancy | State 1 (VIS) | 24.8344 | 26.2563 | 12.1937 | 11.5368 | -1.0566 | 0.2915 |
|  | State 2 (SC) | 26.3642 | 22.9938 | 16.4458 | 15.2478 | 1.8784 | 0.0618 |
|  | State 3 (SM) | 10.8278 | 21.4625 | 17.3154 | 27.5117 | -4.0524 | **6.42e-05\*** |
|  | State 4 (CB) | 48.7682 | 38.6750 | 26.3351 | 23.1616 | 3.5939 | **3078e-04\*** |
|  | State 5 (DMN) | 26.2053 | 27.6125 | 12.4725 | 11.7093 | -1.0262 | 0.3056 |
| Dwell-time | State 1 (VIS) | 16.1788 | 17.8125 | 9.2413 | 9.2186 | -1.5601 | 0.1198 |
|  | State 2 (SC) | 24.5629 | 21.4563 | 16.4534 | 14.9338 | 1.7425 | 0.0824 |
|  | State 3 (SM) | 10.6878 | 21.2688 | 17.1799 | 27.4076 | -4.0513 | **6.44e-05\*** |
|  | State 4 (CB) | 46.2583 | 36.4125 | 26.6542 | 23.2226 | 3.4785 | **5.76e-04\*** |
|  | State 5 (DMN) | 17.9868 | 19.6313 | 9.6540 | 10.0186 | -1.4725 | 0.1419 |

\* Significant at $p < 0.05$ FDR corrected

**Table 2.**
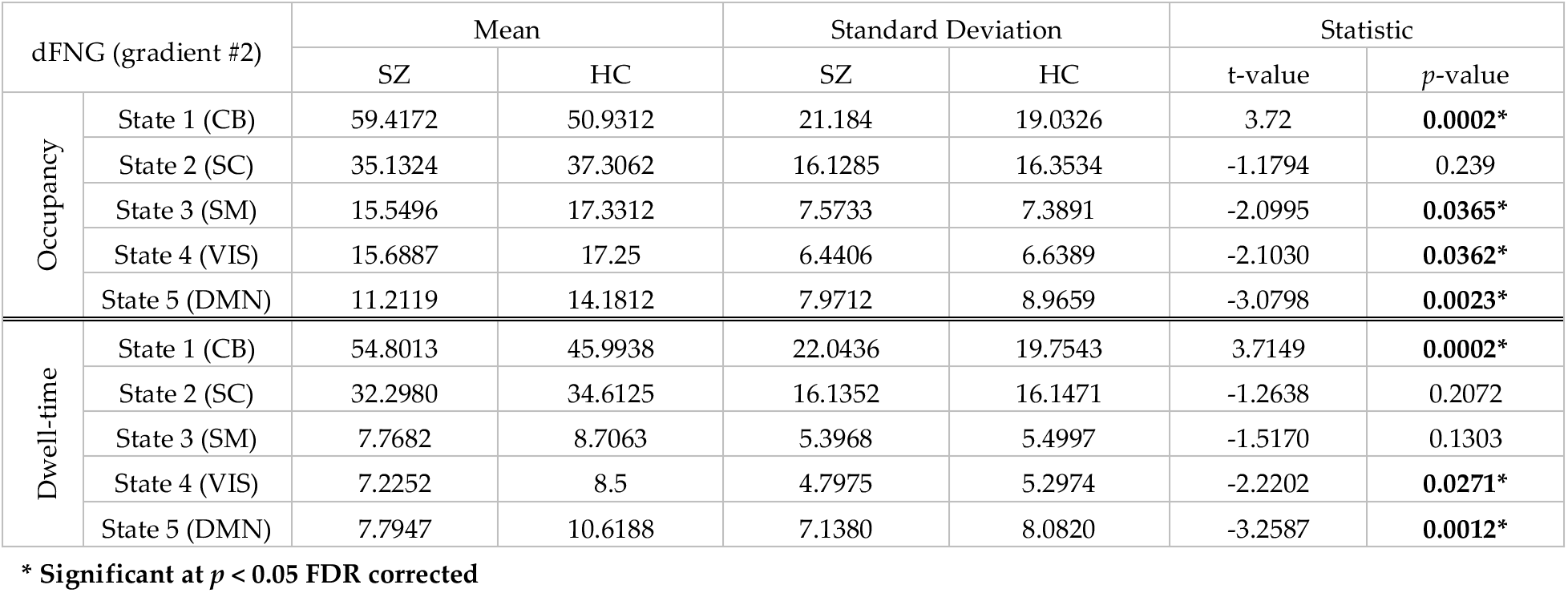
Statistical Results associated with dFNG based on gradient #2.

| dFNG (gradient #2) |  | Mean |  | Standard Deviation |  | Statistic |  |
| --- | --- | --- | --- | --- | --- | --- | --- |
|  |  | SZ | HC | SZ | HC | t-value | p-value |
| Occupancy | State 1 (CB) | 59.4172 | 50.9312 | 21.184 | 19.0326 | 3.72 | **0.0002\*** |
|  | State 2 (SC) | 35.1324 | 37.3062 | 16.1285 | 16.3534 | -1.1794 | 0.239 |
|  | State 3 (SM) | 15.5496 | 17.3312 | 7.5733 | 7.3891 | -2.0995 | **0.0365\*** |
|  | State 4 (VIS) | 15.6887 | 17.25 | 6.4406 | 6.6389 | -2.1030 | **0.0362\*** |
|  | State 5 (DMN) | 11.2119 | 14.1812 | 7.9712 | 8.9659 | -3.0798 | **0.0023\*** |
| Dwell-time | State 1 (CB) | 54.8013 | 45.9938 | 22.0436 | 19.7543 | 3.7149 | **0.0002\*** |
|  | State 2 (SC) | 32.2980 | 34.6125 | 16.1352 | 16.1471 | -1.2638 | 0.2072 |
|  | State 3 (SM) | 7.7682 | 8.7063 | 5.3968 | 5.4997 | -1.5170 | 0.1303 |
|  | State 4 (VIS) | 7.2252 | 8.5 | 4.7975 | 5.2974 | -2.2202 | **0.0271\*** |
|  | State 5 (DMN) | 7.7947 | 10.6188 | 7.1380 | 8.0820 | -3.2587 | **0.0012\*** |

\* Significant at $p < 0.05$ FDR corrected

Figures 7 and 8 show a surface-based visualization of the spatial maps associated with each state of dFNG based on first and second gradient respectively. We also provide a montage view of the 3D spatial maps based on the first and second gradient associated with each state in the appendix.

**Figure 7.**
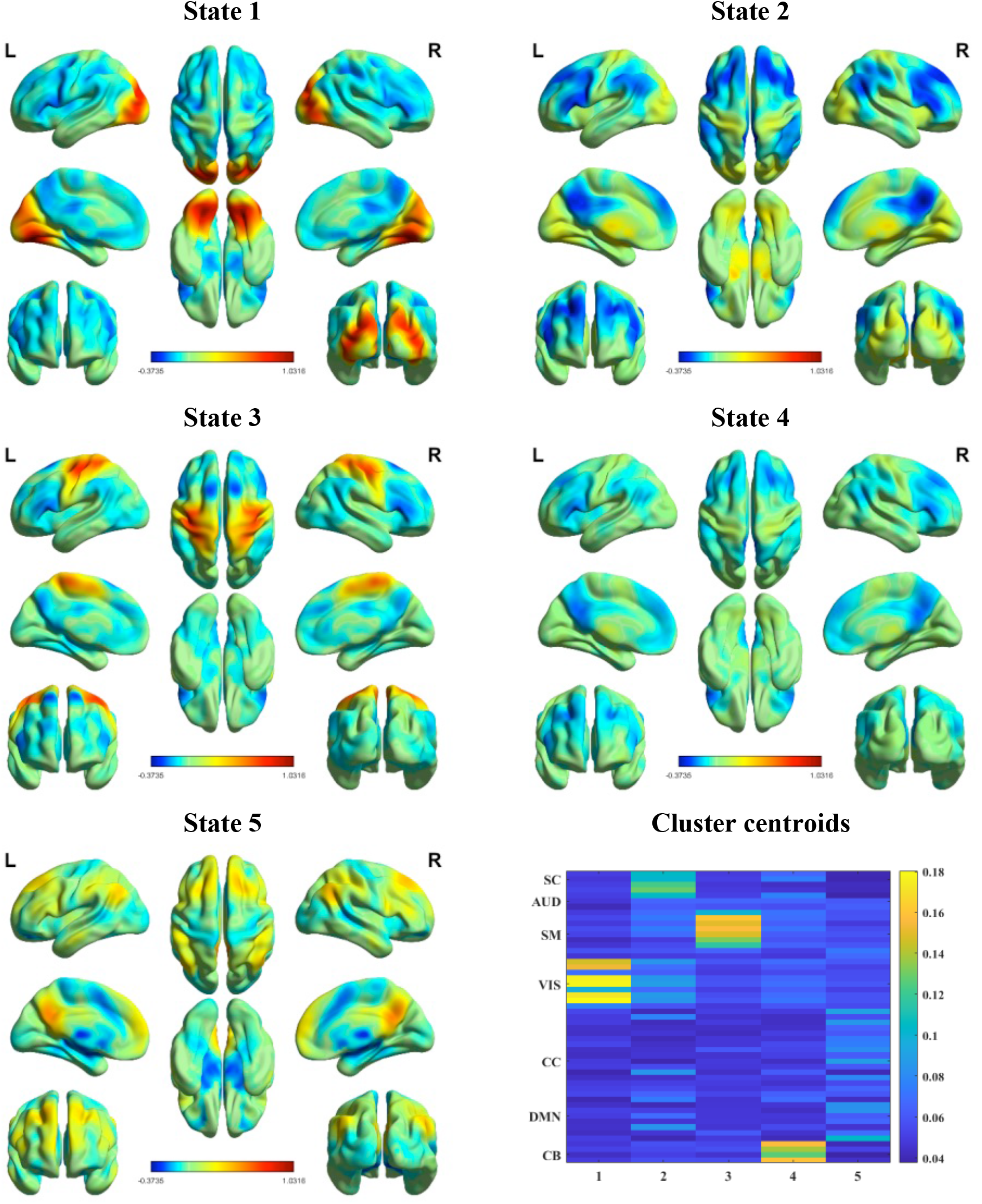
The 3D spatial maps associated with each state based on gradient #1. A spatial map of the dFNGs was created by thresholding and normalizing each component map, followed by using the normalized cluster centroids obtained from gradient #1 k-means clustering as the weight for the component maps to create spatial maps.

**Figure 8.**
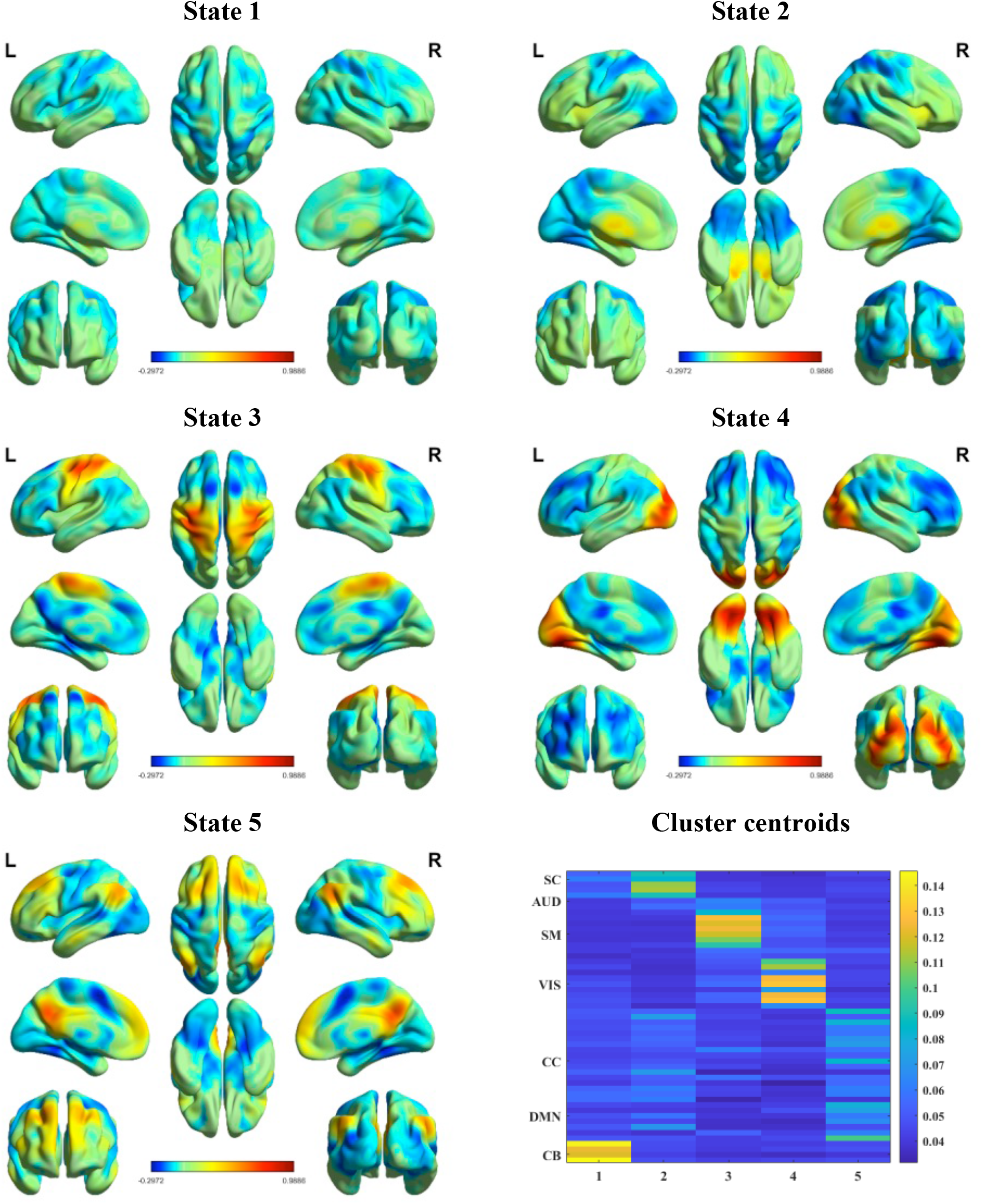
The 3D spatial maps associated with each state based on gradient #2. A spatial map for dFNG was created by thresholding and normalizing each component map, followed by using the normalized cluster centroids obtained from gradient #2 k-means clustering as the weight for the component maps to create spatial maps.

### 3.3. Inter-Component Ordering Synchrony Analysis

Regarding the sort order analysis, no significant differences were found in periodicity between SZ and HC. Periodicity, in this context, refers to the regularity or pattern of sort order across time. Upon closer examination of the data, a notable symmetric pattern emerged in the sort orders of both groups. This led to further investigation into the periodicity of these patterns, which revealed that despite the observed symmetry, there were no statistically significant differences in how participants with schizophrenia (SZ) and healthy controls (HC) exhibited periodicity in their sorting behaviors.

However, the inter-component ordering synchrony analysis, which used to follow the trajectories of sorting orders for each independent component and subjects-capturing the changing order of these components over time-showed significant differences between groups. Initially, distinct sort order patterns emerged over time for each component. Subsequently, cross-correlation analyses were conducted to compare the sort order profiles of each component within each subject, between the two groups. This method allowed for the examination of the degree of similarity in the organization of items over time, providing insights into the underlying cognitive processes and potential differences between subjects diagnosed with schizophrenia and healthy controls. Figure 9 revealed the dynamic gradient ordering vectors associated with one of the healthy controls and the inter-component ordering synchrony plot for component #53.

**Figure 9.**
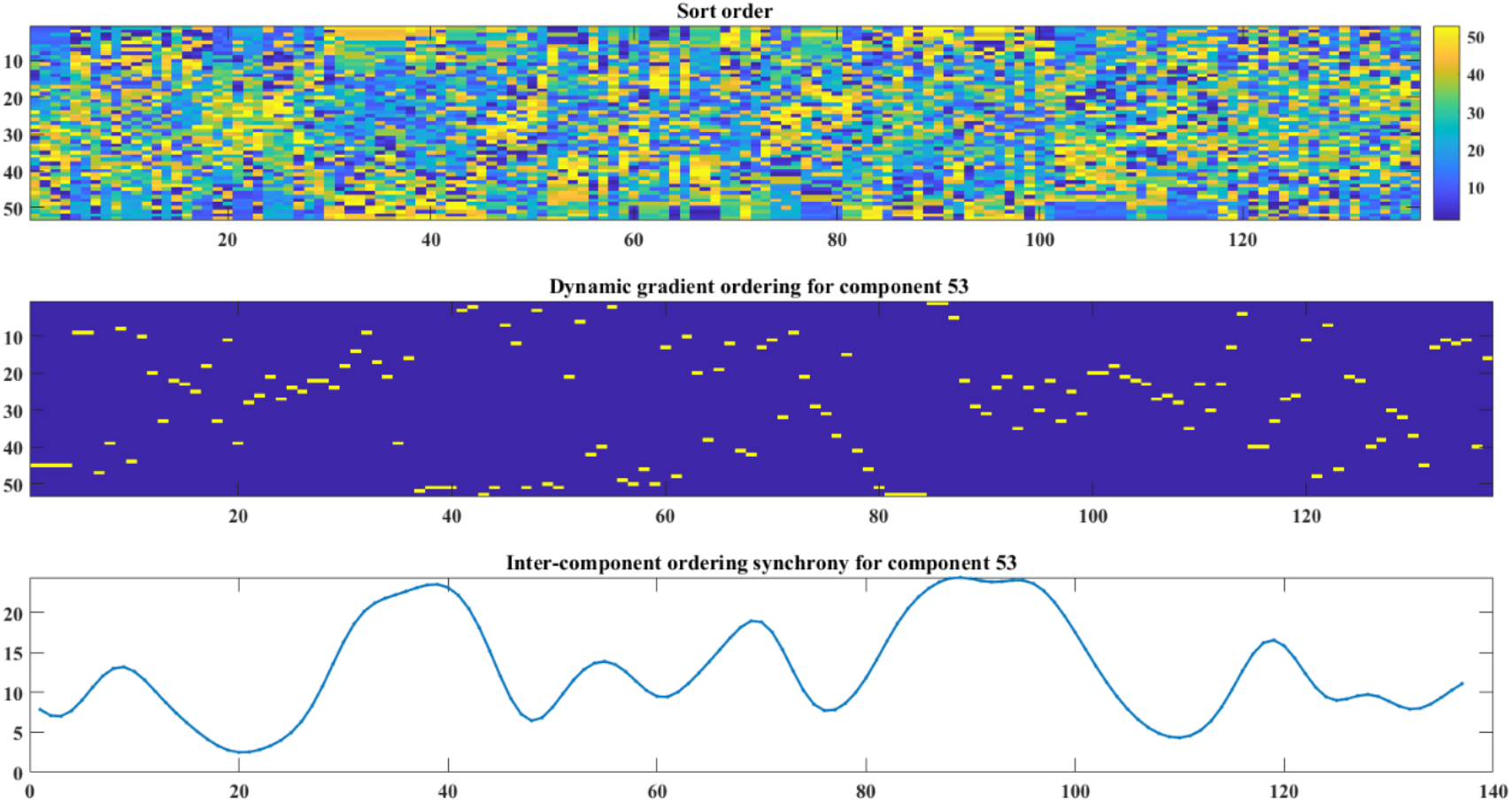
(top) Dynamic gradient ordering vectors for a healthy subject (bottom), dynamic gradient ordering associated with component #53 (middle), and the associated inter-component ordering synchrony plot for component #53.

Figure 10 provides information about the difference between schizophrenia patients (SZ) and healthy controls (HC) in terms of inter-component ordering synchrony. The middle components (DMN/CC/SM) showed significantly higher values in healthy controls in comparison with patients; however, the cross correlation between the end components (SC/CB) were significantly lower in schizophrenia patients.

**Figure 10.**
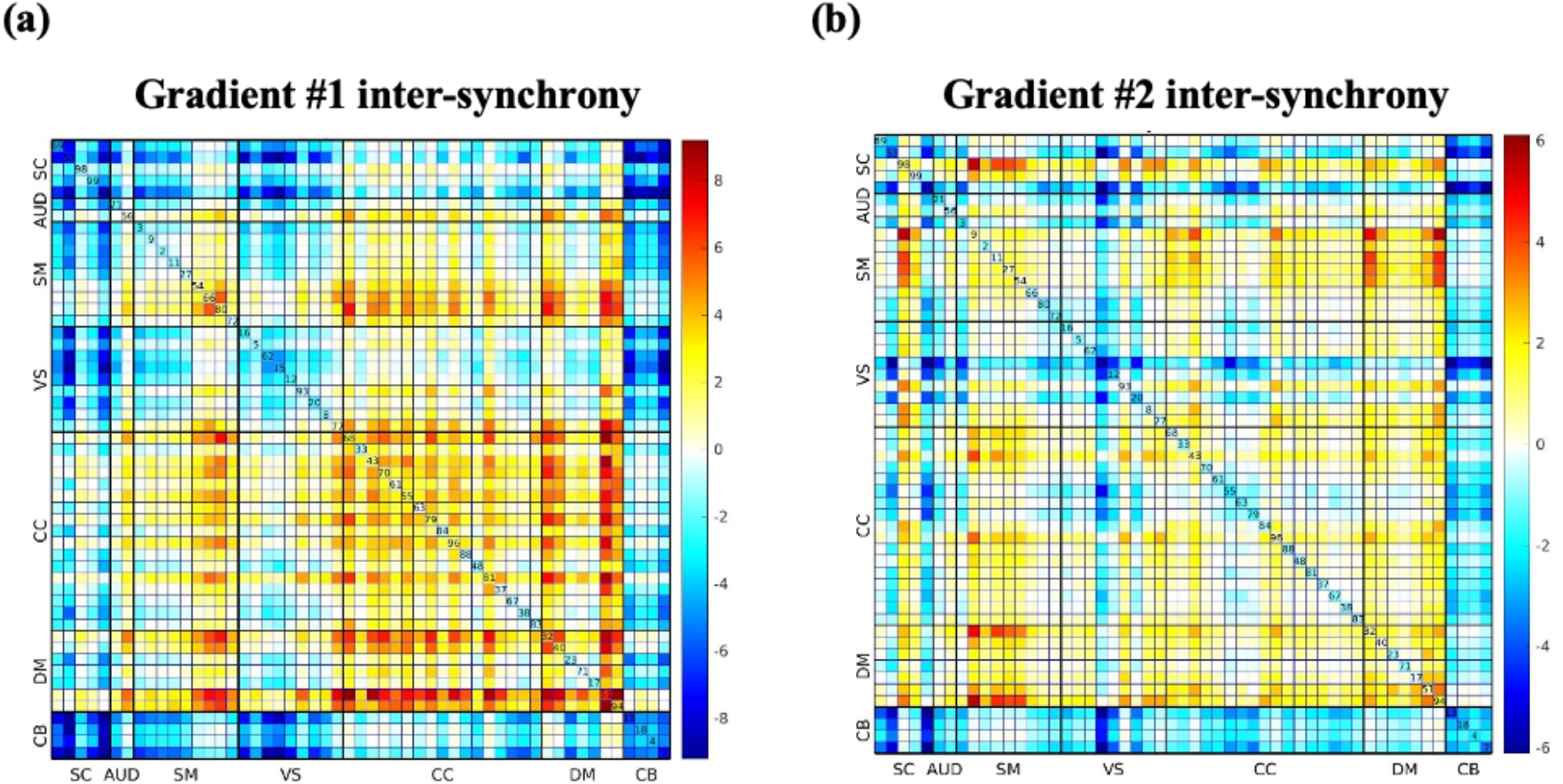
The group difference map associated with inter-component ordering synchrony plot defined as − log!“(*p*value) × sign (tstatistics) for a) gradient #1 and b) gradient #2. After demeaning and smoothing the index order to create inter-component ordering synchrony plot associated with each component for each subject, the cross correlation across all lags is computed, followed by taking the maximum lag for each subject and comparing between patients and healthy controls. The DMN/CC/SM showed significant higher value in healthy controls in comparison with patients, however, the cross correlation between end components (SC/CB) were significantly lower in schizophrenia patients.

The dwell time/occupancy results for the first and second gradient is provided in Table 3 and 4. After computing the gradients followed by k-means clustering, the dwell time and occupancy associated with each state is computed. A two-sample t-test is applied to investigate the group differences. All significant results are shown in bold, with those survived after FDR correction are identified with an asterisk.

**Table 3.** Statistical Results associated with unsigned gradient #1.

| gradient #1 |  | Mean |  | Standard Deviation |  | Statistic |  |
| --- | --- | --- | --- | --- | --- | --- | --- |
|  |  | SZ | HC | SZ | HC | t-value | p-value |
| Occupancy | State 1 (VIS) | 40.8278 | 45.0937 | 32.6271 | 28.5149 | -1.2295 | 0.219 |
|  | State 2 (SC) | 15.2582 | 21.8187 | 15.1078 | 19.2438 | -3.3309 | **0.0009\*** |
|  | State 3 (SM) | 26.8145 | 22.8125 | 23.9792 | 17.7112 | 1.6806 | 0.093 |
|  | State 4 (CB) | 12.298 | 19.25 | 15.4578 | 19.8988 | -3.4267 | **0.0006\*** |
|  | State 5 (DMN) | 41.8013 | 28.025 | 31.5480 | 24.7236 | 4.2992 | **2.3 e-05\*** |
| Dwell-time | State 1 (VIS) | 39.9337 | 44.0375 | 32.5245 | 28.600 | -1.1832 | 0.237 |
|  | State 2 (SC) | 14.1589 | 20.5875 | 14.9381 | 19.0726 | -3.2961 | **0.001\*** |
|  | State 3 (SM) | 25.8543 | 21.9187 | 23.7532 | 17.5531 | 1.6681 | 0.096 |
|  | State 4 (CB) | 11.6622 | 18.3937 | 15.24680 | 19.6023 | -3.3667 | **0.0008\*** |
|  | State 5 (DMN) | 40.5761 | 26.9937 | 31.6253 | 24.6069 | 4.2403 | **2.95 e-05\*** |

\* Significant at $p < 0.05$ FDR corrected

**Table 4.**
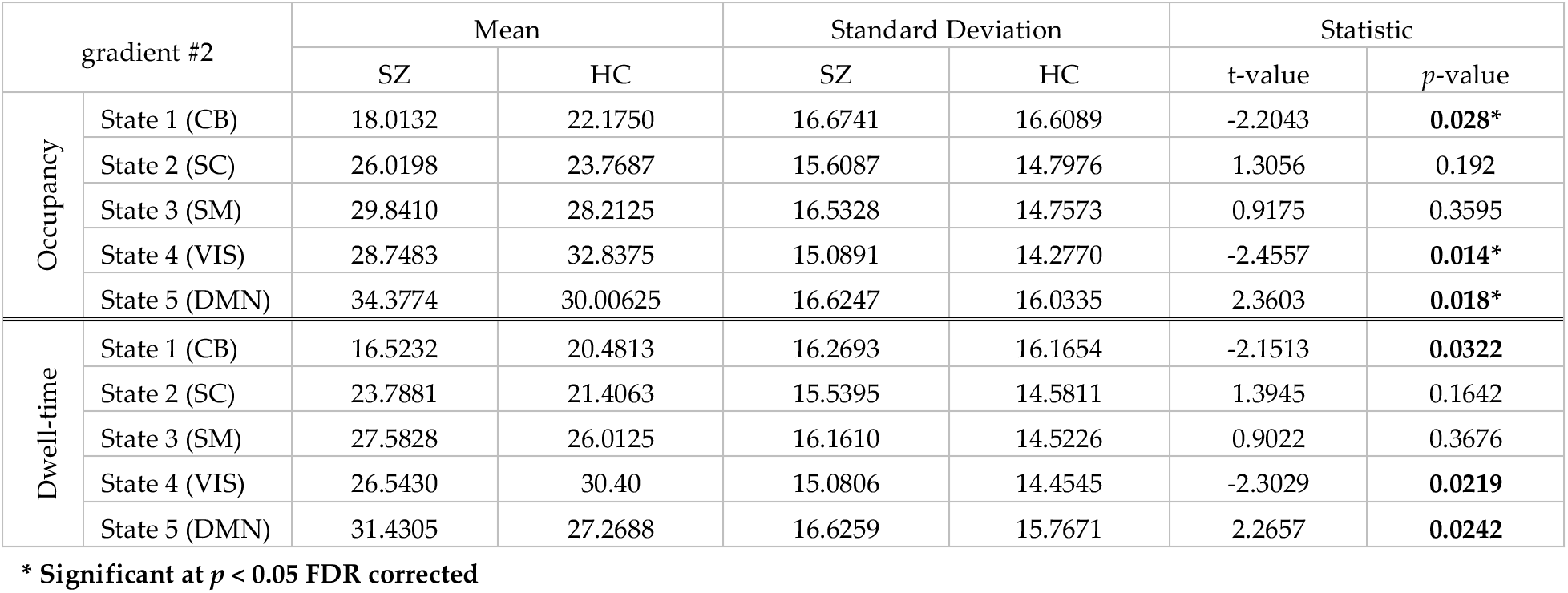
Statistical Results associated with unsigned gradient #2.

| gradient #2 |  | Mean |  | Standard Deviation |  | Statistic |  |
| --- | --- | --- | --- | --- | --- | --- | --- |
|  |  | SZ | HC | SZ | HC | t-value | p-value |
| Occupancy | State 1 (CB) | 18.0132 | 22.1750 | 16.6741 | 16.6089 | -2.2043 | **0.028\*** |
|  | State 2 (SC) | 26.0198 | 23.7687 | 15.6087 | 14.7976 | 1.3056 | 0.192 |
|  | State 3 (SM) | 29.8410 | 28.2125 | 16.5328 | 14.7573 | 0.9175 | 0.3595 |
|  | State 4 (VIS) | 28.7483 | 32.8375 | 15.0891 | 14.2770 | -2.4557 | **0.014\*** |
|  | State 5 (DMN) | 34.3774 | 30.00625 | 16.6247 | 16.0335 | 2.3603 | **0.018\*** |
| Dwell-time | State 1 (CB) | 16.5232 | 20.4813 | 16.2693 | 16.1654 | -2.1513 | **0.0322** |
|  | State 2 (SC) | 23.7881 | 21.4063 | 15.5395 | 14.5811 | 1.3945 | 0.1642 |
|  | State 3 (SM) | 27.5828 | 26.0125 | 16.1610 | 14.5226 | 0.9022 | 0.3676 |
|  | State 4 (VIS) | 26.5430 | 30.40 | 15.0806 | 14.4545 | -2.3029 | **0.0219** |
|  | State 5 (DMN) | 31.4305 | 27.2688 | 16.6259 | 15.7671 | 2.2657 | **0.0242** |

\* Significant at $p < 0.05$ FDR corrected

The cross correlation between the cluster centroids (states) were also computed for HC and SZ. Figure 11 provide information about the difference in correlation between the gradient centroids. The HC-SZ plot showed that the connectivity between the second centroid (SC) with SM and CB is positive in controls and negative in patients for both the third centroid (SM) and the fourth centroid (CB).

**Figure 11.**
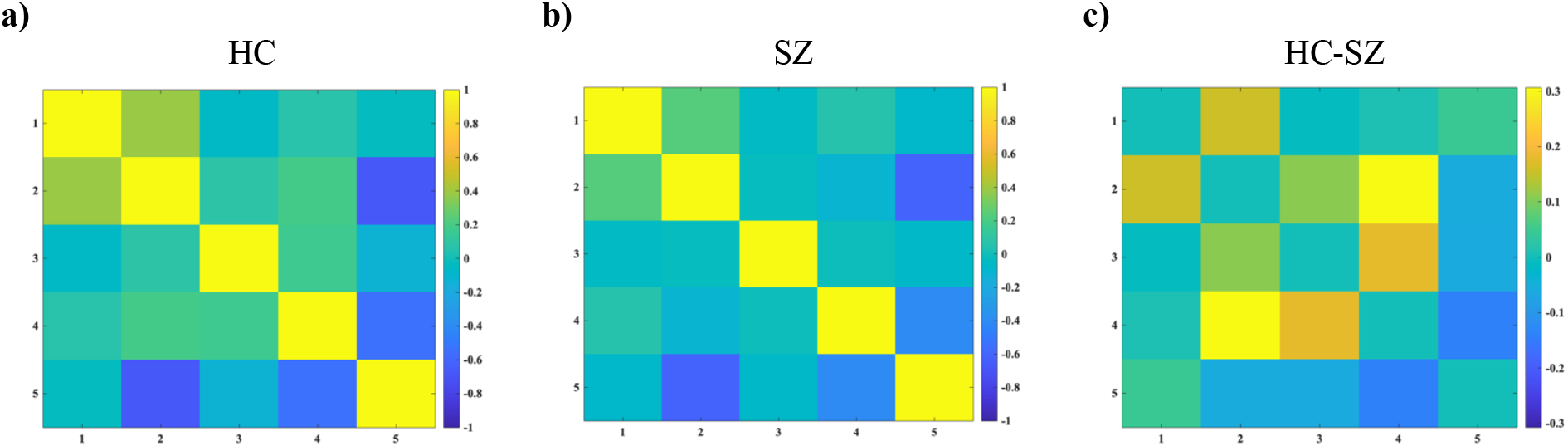
The cross correlation between cluster centroids for a) healthy controls (HC), b) patients with schizophrenia (SZ), and c) healthy controls – patients (HC – SZ). The HC-SZ map is representative of the positive connectivity between the second centroid with SM and CB in controls and negative connectivity in patients for both the third centroid (SM) and the fourth centroid (CB).

## 4. Discussion

Recent empirical studies report on the temporal reconfiguration of functional connectivity and dynamic properties of the brain [9],[37]. Their findings suggest that the spatial and temporal properties of neural activity interact on several spatiotemporal scales [5],[9]. This has encouraged the development of new approaches focused on temporally static spatial topography (e.g., spatial cortical gradients) of brain connectivity [15],[38],[39]. Gradient-based approaches provide an organizational framework for capturing the complex large-scale structural and functional organization of the brain [39],[40]; However, brain activity is ever changing and the functional topography may change accordingly [41]. Furthermore, there has not yet been a focus on studying the degree to which these gradients might fluctuate over a short time frame, and how this might provide insights into the spatio-temporal behavior of fMRI data and its application to understand the pathophysiology of schizophrenia.

This study highlights the potential of dFNGs as a new method for understanding the spatiotemporal dynamics of brain and dysfunction. We investigate the smooth transitions caused by dFNGs from ICNs, as well as sFNG. In parallel with the proposed approach, the effects of cortical gradient were also studied in a group of 151 individuals with schizophrenia (SZ) in comparison with age and gender-matched healthy controls (HC). Our main findings are that: 1) multidimensional interactions of the first two gradients are clustered along three networks of CC/ DMN, SC/ AUD/ SM/ CB and VIS in both SZ (Figure 4a) and HC (Figure 4b); 2) that sFNG differ in the SC, CB, and DMN between SZ and HC; 3) that occupancy of state 4 (CB) is higher in SZ compared to HC based on the first cortical gradient, 4) that occupancy of state 1 (CB) is higher in SZ comparison to HC based on the second cortical gradient; 5) that compared to HC, SZ shift more between the end (SC/ SM) and middle components (CC/ DMN) based on the inter-component ordering synchrony analysis, and 6) there is positive connectivity between the second centroid with SM and CB region in HC, and negative connectivity for both the third (SM) and forth centroids (CB) in SZ based on the gradient centroids cross correlation analysis.

These findings suggest that sFNG and dFNG can aid in characterizing differences in the global organization of functional brain networks, and dynamic changes in brain connectivity between SZ and HC, respectively. Dynamic analyses have revealed fluctuations in gradient strength and variability over time, reflecting the flexible reconfiguration of brain networks. In addition to the emerging consensus that gradients may represent important patterns of intrinsic brain organization [21],[40], it remains to be investigated how far these patterns constrain state-to-state variation in brain function. In line with previous task-evoked studies, the magnitude of regional activity is high in unimodal networks (e.g., primary sensorimotor regions), but low in transmodal regions (e.g., DMN) in healthy controls [40]. Also pointing to hierarchy-dependent shifts in localized vs distributed processing. Recent advances in neuroimaging methods enable us to use cortical gradients as a dimensionality reduction method. Gradient approaches have been able to find the main axes of variance in the data through embedding techniques. The original dimensions of the data are replaced by a set of new dimensions, so that most of the variance in the data is captured by just a few of these dimensions [40],[42]. Each dimension is a large-scale cortical gradient. To put it simply, each dimension can be representative of one aspect or network of cortical organization. In line with our results regarding the multidimensional interaction between the computed gradients which seems to be aligned along three domains of VIS, SC/AUD/SM/CB and CC/DMN networks. Furthermore, utilizing dynamic rs-fMRI analysis, Yousefi and colleagues demonstrate how intrinsic functional activity propagates along macroscale functional gradients [43], suggesting that these axes may play a role in constraining functional dynamics.

The observed differences between SZ and HC in SC, CB, and DMN extend recent reports using ICA [20],[22],[44]. By investigating the whole brain functional connectivity, stronger connectivity between the thalamus and sensory networks (auditory, motor and visual), as well as weaker connectivity between sensory networks were reported [20]. Using seed-based connectivity, Woodward and colleagues also reported stronger functional connectivity between the subcortical and somatosensory regions in patients with schizophrenia compared to healthy controls [45]. Our sFNG results also suggest the weaker connection between SC and CB ICNs in patients. This, apparently novel, finding is present in data. The identification of this group difference, along with connectivity differences related to subcortical areas, speaks to the strength of our whole-brain, data-driven approach, which is not limited by the selection of any specific region of interest.

Using a dynamic analysis based on sliding windows and k-means clustering of cortical gradients, we identified five different states (Figure 7 and 8). We found that SZ, compared to HC, spend significantly longer duration in state 2 and 4, as well as 1 and 4, based on gradient 1 and gradient 2 respectively, which are associated with SC, CB and VIS. These findings are consistent with those from prior studies which have identified reproducible neural states in a data-driven manner and demonstrated that the strength of connectivity within those states differed between SZs and HCs [44].

### 4.1. Limitations

While the presented study offers valuable insights into brain network dynamics using a novel approach of dynamic functional network connectivity gradient analysis, several limitations should be acknowledged. First, the generalizability of the findings may be constrained by the specific dataset utilized, consisting of 151 schizophrenia patients and 160 age and gender-matched healthy controls. Larger and more diverse samples could provide a broader representation of the population and enhance the robustness of the results. Furthermore, due to the use of the cross-sectional research design, we did not establish the developmental trajectories of altered cortical hierarchy in schizophrenia. Future longitudinal studies may evaluate the development of cortical hierarchy in schizophrenia across time.

In sum, while the study advances the field by introducing a novel approach to characterizing brain network modulation, these limitations underscore the need for further research. Addressing these challenges could enhance the reliability, validity, and clinical relevance of dFNG analyses in the context of mental disorders and beyond.

## 5. Conclusions

The present study investigated the static and dynamic functional network connectivity using spatial gradients rather than assuming fixed spatial maps for evaluating the transient changes in coupling among independent component time courses. A summary of the sFNG, the dFNG and its reordering properties, and the dynamics of the gradients themselves were evaluated. This approach was applied to a dataset of individuals with schizophrenia and healthy controls to investigate group effects of these findings as well as the ability to detect differences between individuals with a clinical diagnosis and healthy controls. Regarding the sFNG analysis the gradients interaction showed the gradient values are relatively clustered along three networks of (CC/ DMN), (SC/ AUD/ SM/ CB) and (VIS) for both schizophrenia patients (SZ) and healthy controls (HC). Significant differences in the sFNGs were observed in SC and CB regions. dFNG analysis suggests that SZ, compared to controls, spend a longer duration in cerebellar network (CB). Furthermore, the ordering index cross-correlation of each component line plot was representative of the patients shifting between the end (SC/ SM) and middle components (CC/ DMN), and the cross-correlation between the gradient centroids of healthy controls showed aberrant pattern in connectivity pattern of second centroids with DMN and SC. Finally, by employing the dFNG from ICA, we leverage both higher order statistics and spatial smoothness, to provide a more complete spatiotemporal summary of the resting fMRI data.

## Acknowledgement

This work was supported by the National Institutes of Health (NIH) grant number R01MH123610 and National Science Foundation (NSF) grant number 2112455 to Vince Calhoun.

## Competing Interest

The authors declare no competing interests.

## Appendix

**Figure A1.**
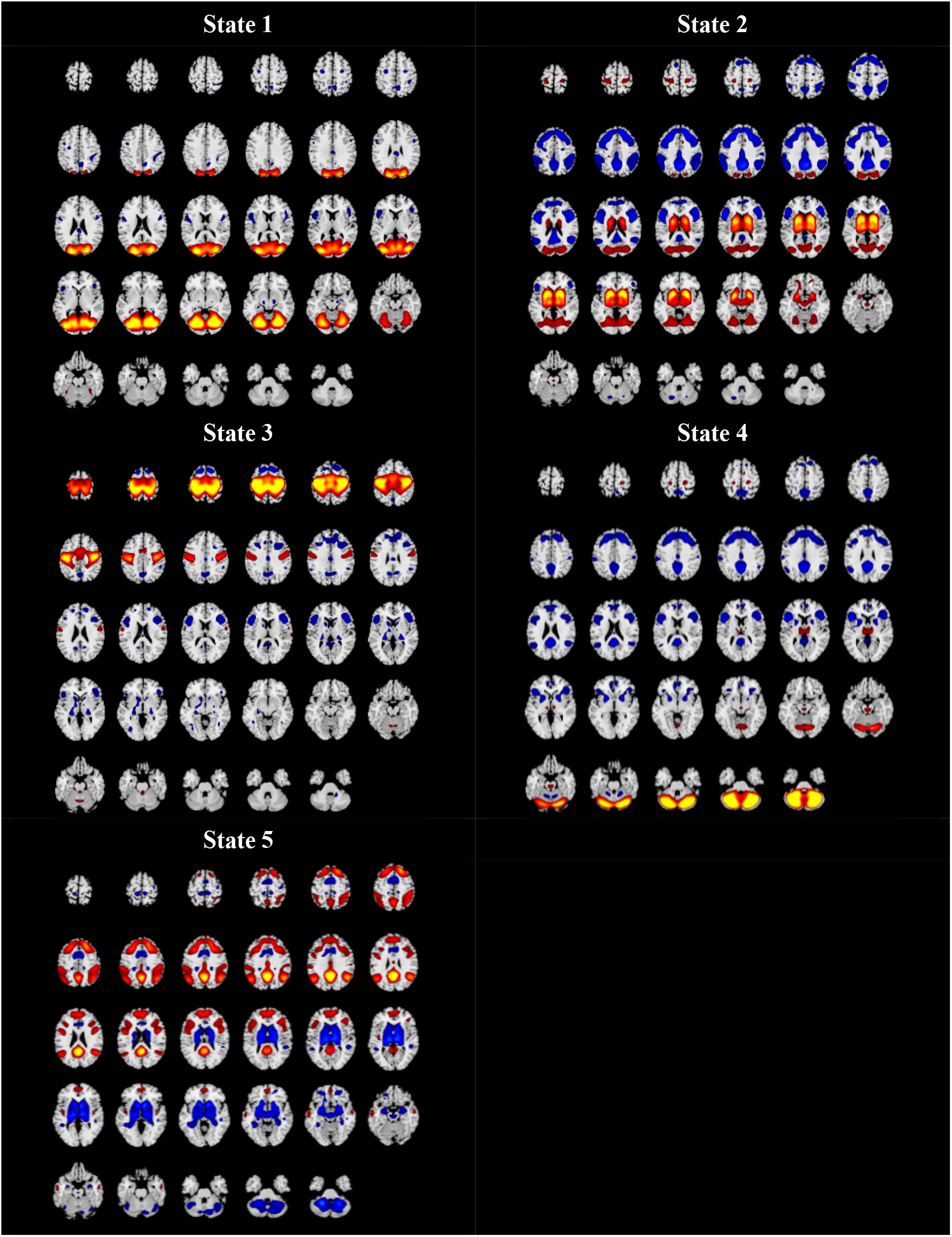
The 2D Spatial maps associated with each state based on the gradient #1. A spatial map of the dFNG was created by thresholding and normalizing each component map, followed by using the normalized cluster centroids obtained from the gradient #1 as the weight for the component maps to create spatial maps.

**Figure A2.**
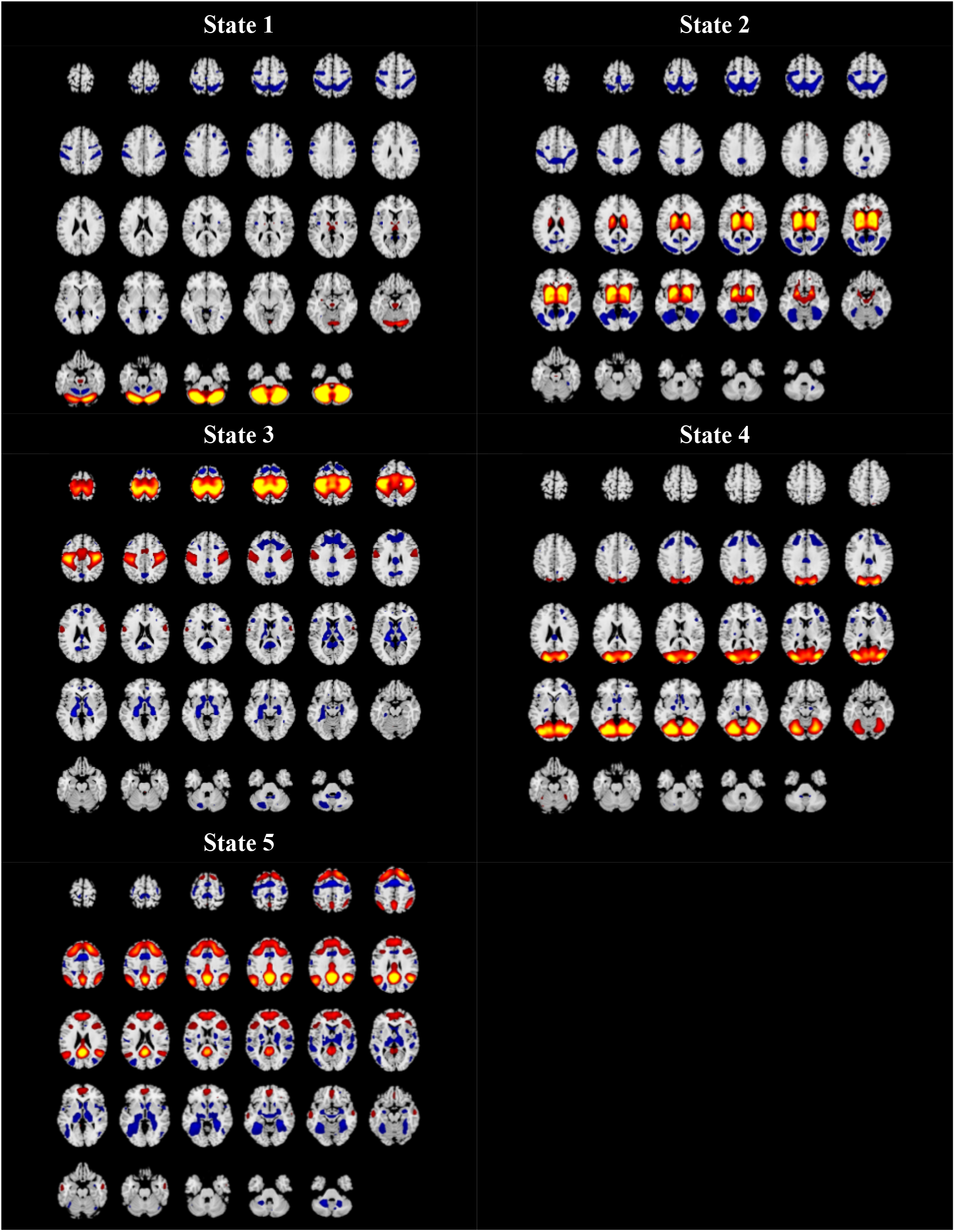
The 2D Spatial maps associated with each state based on the gradient #2. A spatial map of the dFNGs was created by thresholding and normalizing each component map, followed by using the normalized cluster centroids obtained from the gradient #1 as the weight for the component maps to create spatial maps.

## Notes

### Competing Interest Statement

The authors have declared no competing interest.

### Summary of Updates

Another Co-author is added as well as some tables which are updated on the result part.

